# Information-based planning for optimal intermittent control

**DOI:** 10.1101/2025.03.05.641580

**Authors:** Atsushi Takagi, Dorian Verdel, Etienne Burdet

## Abstract

Mammalian motor control is inherently discrete, with movement corrections occurring at rates determined by task demands and the quality of sensory information. While several models have been proposed to explain this discreteness, it remains unclear when a new movement should be initiated and how long it should last. To address this gap, we introduce the Information Predictive Control (IPC) framework, which combines model predictive control with information theory. IPC triggers corrections only when unexpected deviations arise and when corrective actions are likely to succeed. By quantifying “surprise” relative to predicted internal and external states, IPC generates successful movements while robustly integrating sensorimotor noise, task constraints, and target variability. Simulations show that IPC reproduces human-like behavior in discrete reaching, continuous target tracking, and adaptive planning under uncertainty, while dynamically adjusting the planning horizon in complex, unpredictable environments.

## 1 Introduction

Mammals face the fundamental challenge of planning movement in a dynamic and unpredictable environment. For instance, they must continuously replan their actions to intercept an agile prey and interact efficiently with a dynamic environment. Early attempts to model motor planning mechanisms relied on deterministic feedforward control, optimizing energy or motion smoothness to predict the average trajectory across repeated trials [1–7]. More recent approaches have accounted for internal or external noise [8–10] to predict movement trajectories and human impedance. While these open-loop control models perform well when internal and external dynamics change slowly, allowing the brain to build reliable internal models [11], they are not designed to react to sudden changes, such as for catching a fly, or recovering from a stumble after hitting the sidewalk.

In such dynamic situations, the ability to react to task or environmental changes by replanning movement is essential [12]. The continuous correction of deviations from a goal and the associated effort has been modeled using Optimal Feedback Control (OFC) [13, 14], where the corrective strategy adopted by humans may adapt within single [15] or between multiple trials [16, 17]. How OFC is adapted in dynamic environments has been recently summarized in [18]. However, OFC yields a servo mechanism responding to any target movement without distinguishing whether the target is unreachable, due for example to sensory noise or rapid oscillations. Moreover, originally developed for aeronautics, the basic OFC mechanism fails to predict key features of mammalian motor control, particularly the submovements observed in cats [19], mice [20], humans [21–23] and non-human primates [24]. These submovements suggest that mammals’ sensorimotor control may rely on an intermittent strategy.

Several mechanisms have been proposed to explain movement intermittency in humans. It has been attributed to biological constraints imposing a *fixed-rate control* (FRC) frequency [21], then formalized in an optimal control framework [25]. It may also arise due to motion replanning, either based on (i) a task-error threshold, which is not directly applicable to reaching movements as the threshold would always be exceeded, triggering repeated replanning events [26, 27], or on (ii) an error between prediction of upcoming movement and the actual trajectory, resulting in a *prediction error threshold control* (PETC) [28]. However, these explanations do not account for observed variations in control frequency with the sensory feedback quality, i.e., movement intermittency decreasing when tracking a target with noisy vision [29]. Furthermore, existing models overlook how motor noise and other sources of deviation influence the replanning rate. Current models also do not adapt the *planning horizon* to evolving environments [30]. A comprehensive framework is needed to integrate these factors and explain how movements are dynamically replanned with environmental changes.

The paper introduces *information-based predictive control* (IPC) as a framework incorporating sensory noise, motor noise, task-related constraints, and environmental unpredictability into motor planning. IPC models internal and external states to predict upcoming conditions, guiding optimal actions through a model predictive control scheme [31, 32] with an adaptive planning horizon. We propose that the central nervous system (CNS) monitors deviations between predicted and actual states, triggering movement replanning when discrepancies exceed a perceptual threshold quantified using Shannon entropy [33] (Fig. 1). Through simulations and comparisons with the FRC and PETC models of intermittent control, we show how IPC generates human-like responses in diverse motor tasks, including arm reaching in a force field, planning under motor noise, continuous target tracking with an adaptive horizon, and hunting prey with varying agility.

**Figure 1:**
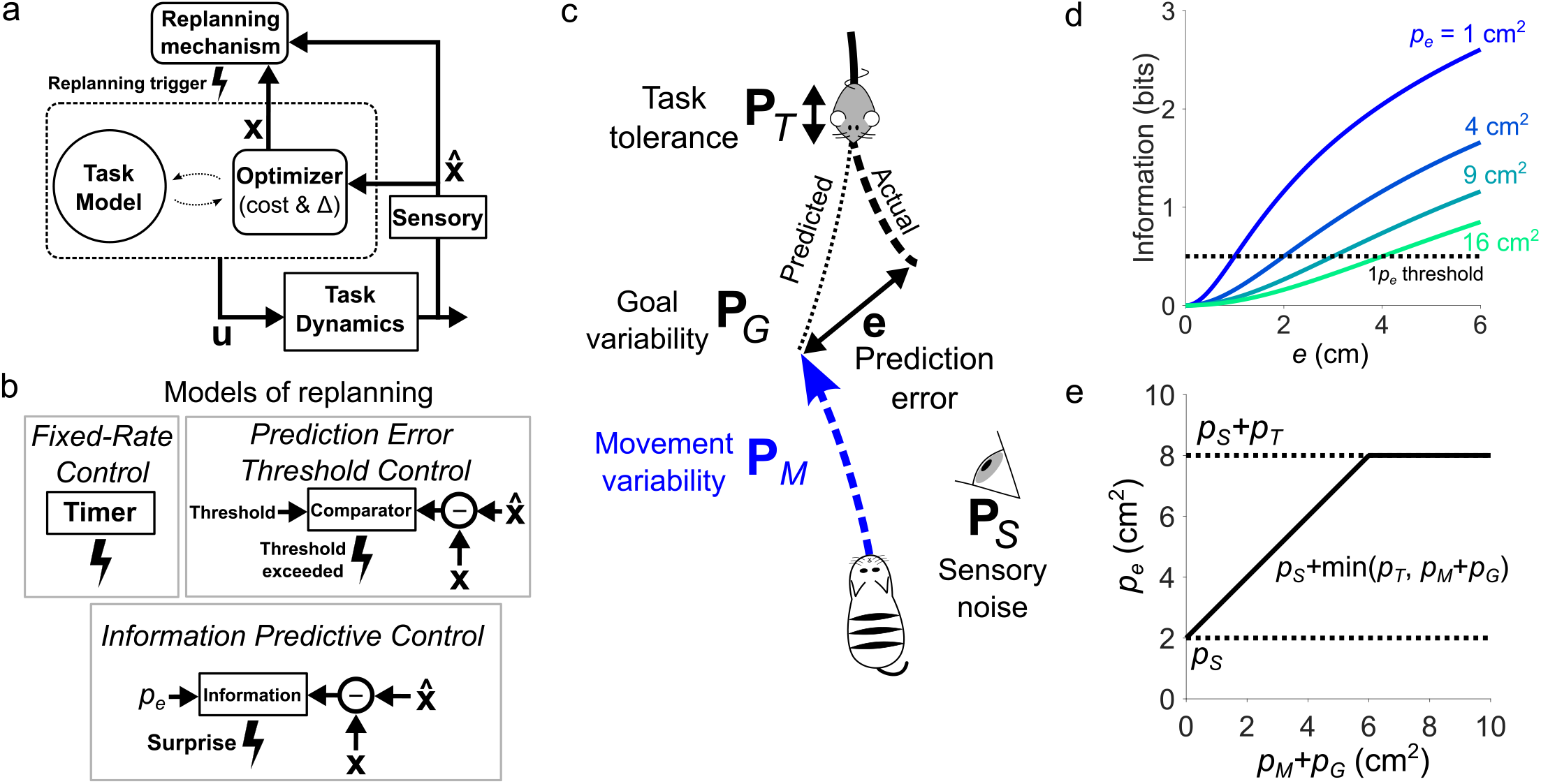
Investigated models for planning and replanning movements. **(a)** General concept of an intermittent controller: after an initial planning, based on the priors of the CNS, a new movement is replanned from the current state over a time horizon Δ when the replanning mechanism is triggered. **(b)** Replanning mechanisms of the three investigated models: (i) FRC replans after a fixed time interval elapses, as in [25]. PETC replans when the prediction error exceeds a threshold, as in [28]. IPC replans after large surprise, measured using Shannon’s information as described below. **(c)** The need for a complete framework such as IPC is clear in a broad range of mammalian activities, such as when a hunter tries to intercept a prey. The hunter must predict the prey’s movement to intercept it. However, if the prey changes its plan to evade capture, or if the hunter’s movement deviates from the planned trajectory due to sensorimotor noises or external events, then a replanning is needed to correct the hunting trajectory. In details, replanning events should be triggered when observed task errors exceed the CNS’ prior knowledge of internal sensory **P**_*S*_ and motor noises **P**_*M*_, variability in the target’s (here the prey) movement **P**_*G*_, and the task tolerance **P**_*T*_. The combination of these elements allows to define a *permissible variance* **P**_*e*_ for a given task. **(d)** In IPC, Shannon’s information depends on the difference between predicted and observed task errors, with a rate of increase inversely proportional to the level of permissible variance. This is because when the permissible variance increases (here with only one-dimensional position deviations for the sake of simplicity, i.e. *p*_*e*_ = **P**_*e*_ ∈ ℝ), IPC deems the same prediction error to be less surprising. **(e)** The permissible variance *p*_*e*_ is capped by the task tolerance *p*_*T*_, which is because a task error exceeding the task tolerance leads to failure. Below this point, *p*_*e*_ is a function of motor and target’s variability *p*_*M*_ + *p*_*G*_. Sensory noise *p*_*S*_ offsets the entire function, which is because errors below the sensory noise level cannot be observed and create surprise.

## 2 Results

### 2.1 Intermittent movement replanning

We start by providing a general principle for the intermittent control of movements (Fig. 1a). Let **x** represents the system’s state and **x**^∗^ the goal’s (or target) state. These states of dimension *N* may include position, velocity, force, etc. We consider three models allowing to trigger replanning events while a movement is executed. First, the FRC model triggers replanning events at a fixed rate (Fig. 1b). In our simulations, the time interval was set to 130 ms, corresponding to a replanning rate of ∽ 8 Hz as observed in [25]. Second, the PETC model triggers replanning events when the magnitude of the *prediction error* between predicted and observed task errors

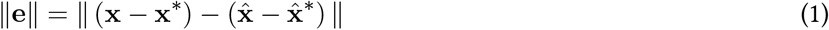

exceeds a threshold [28, 34, 35], where 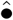 are predicted variables.

Last, in contrast to these previous methods, the IPC model we introduce here replans a movement when the Shannon information *I* [33] generated by the prediction error exceeds a threshold *I*_*T*_ :

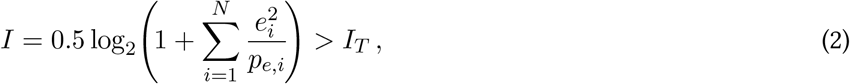

where **p**_*e*_ is the permissible state variance vector. In this study we set *I*_*T*_ = 0.5, meaning that a deviation exceeding one standard deviation of the prediction error is considered sufficiently “surprising” to initiate replanning.

As illustrated in Fig. 1d, when **p**_*e*_ components are small, even minor prediction errors generate high information values, making IPC more sensitive to deviations. The *permissible variance* depends on the internal sensorimotor noise, the target’s variability, and task tolerance as follows (Fig. 1c),

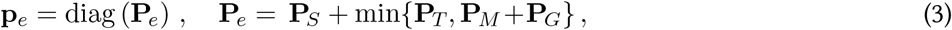

where the minimum is defined component wise and the respective sources of variability are assumed to be independent stochastic variables. Fig. 1e illustrates how these four factors influence the permissible variance. Specifically, an increase in sensory noise reduces the reliability of the prediction error estimate, making IPC less likely to trigger replanning. Similarly, as the internal values attributed to movement and goal variabilities increase, replanning becomes less likely, until their sum exceeds the task-admissible variability. This implies that the internal movement and goal variabilities will set the limit of total admissible variance, as long as their sum remains below the task tolerance. This mechanism allows to implement a task-constraints-aware minimum intervention principle, extending previous works [13,36]. When the motor noise or the target variability exceeds the task tolerance, the likelihood of replanning increases because the prediction error more often exceeds the permissible variance. If IPC predictions are accurate, the prediction errors will remain small enough so as not to trigger replanning (Fig. 1d). In contrast, when sensorimotor noise or unexpected target motion occurs, IPC’s prediction errors grow over time, triggering replanning.

## 2.2 Cost of movement

A common framework is used to simulate movement planning with FRC, PETC, IPC, i.e. only the replanning decision process differed between these three models. We assume that the brain maintains an internal model of the task dynamics 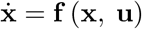 and 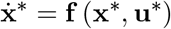, where **u** is the control vector of the system typically including joint torques or muscle activations. This internal model enables the brain to predict the trajectory of the goal state **x**^∗^ and plan optimal actions over a *finite time horizon* Δ depending on the defined system variances. The learning of this internal model by the brain, which incorporates system dynamics and the four sources of variability and could be modeled using Bayesian or active inference [37, 38], is beyond the scope of this study. Using this internal model, the brain plans actions by minimizing a cost function:

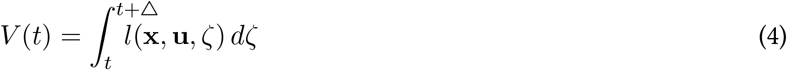

over the horizon Δ where *t* is the current time of replanning and *l*(**x, u**, *?*) a cost function modeling human behavior. In the present paper, we assume that an intermittent controller may generate movements minimizing the distance between own state **x** and the goal state **x**^∗^ with minimal effort. This is implemented with a quadratic function of error to the target and the effort required to reach it, as in continuous approaches to motor control [13, 18, 39] (see Methods for details on cost functions used in simulations).

In the rest of the paper, we investigate the behavior and performance of FRC, PETC, IPC in increasingly complex scenarios, ranging from the seminal reaching under perturbation to the case of a hunter-prey scenario. We thereby investigate the sensitivity of IPC to each of the permissible variance components, either by keeping the others fixed or by setting them to zero. For the sake of simplicity, and without loss of generality, only stochasticity in position will be considered, reducing the permissible variance vector **p**_*e*_ and matrix **P**_*e*_ to position alone. Furthermore, we assume that the variables in Eq. 3 reflect the true variability of the system such that 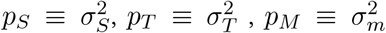 and 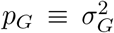 to focus on how these parameters affect the replanning rate and the horizon. As the main contributions of the present paper are IPC’s replanning mechanism and adaptive planning horizon, we did not simulate feedback mechanisms [18], for instance related to reflexes [40, 41], that can generate a corrective force as soon as or just after a mechanical perturbation.

### 2.3 Reaching under perturbations

We first investigate the effects of the task-tolerance **p**_*T*_ = diag(**P**_*T*_) component of the permissible variance, yielding a minimum intervention principle that is a major feature of volitional motor control [13, 36]. This is performed by simulating reaching movements towards disks of various sizes with an unexpected velocity-based force-field (not accounted for in the planning), pushing to the right for a force proportional to the vertical speed [16, 42] (see Methods for details). Fig. 2a illustrates the responses to this unexpected perturbation for each of the three investigated models.

**Figure 2:**
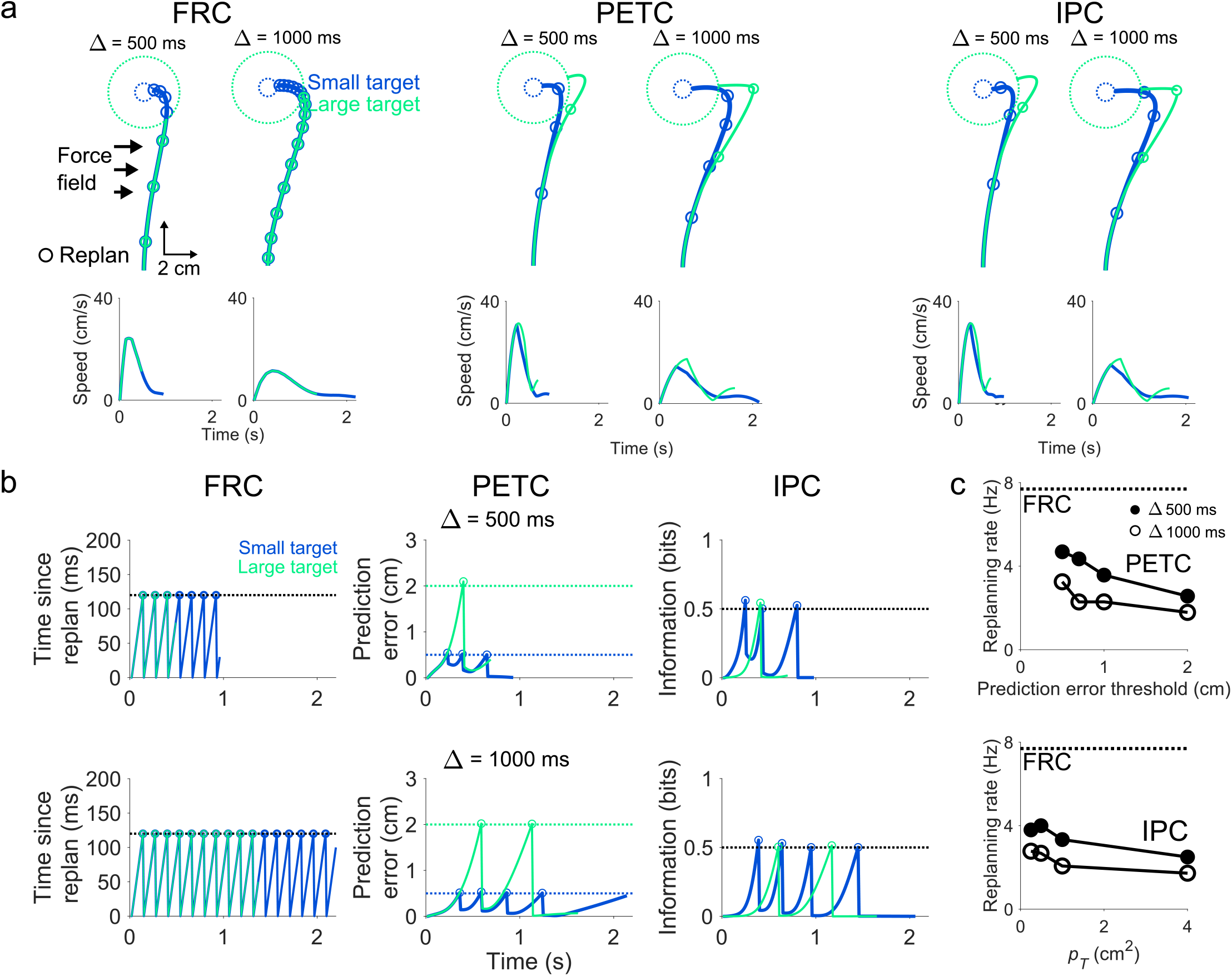
Change in replanning rate with different target sizes. (a) Trajectories and speed profiles when reaching a small (blue) and large target (green) using FRC, PETC, IPC with short or long horizon. (b) Replanning with FRC, PETC, and IPC. FRC plans a new movement after a constant time interval elapses. PETC plans a new movement when the prediction error threshold is exceeded, IPC after a large surprise. With IPC more information is gained for the same prediction error with smaller task tolerance *p*_*T*_. (c) Replanning rate as a function of the prediction error threshold (PETC) and task tolerance (IPC). The replanning rate declines as prediction error threshold and task tolerance increase. FRC’s replanning rate is the same for all target sizes.

For all the controllers, the performed movement deviates from the typical straight line from the start to the target position [2] predicted by the three controllers. This unexpected deviation allows to illustrate the outcomes of the replanning process with each model. We simulate reaching movements towards a small (blue) or large (green) target, and illustrate the controllers’ response for different movement horizon durations Δ by simulating reaching with Δ= 500 ms and Δ= 1000 ms (Fig. 2a).

When the horizon is shorter, all controllers reach the target earlier and with greater speed than with the longer horizon (Fig. 2a). Then we examine how each controller’s movement replanning is affected by the target size, where the circles along the path in Figs. 2a-b indicate the times of replanning. FRC replans with the same constant rate, thus independent on the target size (left panels of Fig. 2b). However, the behavior with PETC and IPC’s change with the target size, where both replan more often when reaching the smaller target while being perturbed laterally (Fig. 2b, middle and right panels). When reaching to four different target sizes (Fig. 2c), the replanning rate of PETC and IPC decreases as the target size increases. PETC and IPC’s increase in replanning rate for smaller target is consistent with observations of human behavior as there is more replanning and submovements when reaching towards smaller targets [43]. Notably, the number of replanning events is higher with FRC than with PETC or IPC, and similar for PETC and IPC, showing that in absence of sensorimotor noise and external uncertainty, predictions of these two models can be close.

Note that FRC, PETC, and IPC also reproduce human-like behavior when learning novel force fields, and produce an after-effect or overcorrection in the opposite direction to the force field when it is abruptly removed after learning (see Fig S1 in the Supplementary Materials).

### 2.4 Optimal horizon adaptation during intermittent control

The previous simulations assumed that the controllers planned the remaining movement at the initial planning or at each replanning event. However, this may result in “over-planning” when their is uncertainty as illustrated in Fig.2a, where the unexpected force-field generates uncertainty, resulting in an unusable plan computed over a long duration for no benefit. Therefore, the horizon should adapt to the uncertainty in the task at hand. This notion will allow us to investigate the effects of a second component of IPC’s permissible variance on replanning, which is the known level of internal motor variability **p**_*M*_ = diag(**P**_*M*_).

It remains unclear how large the horizon should be and according to which principle it should adapt. We propose that if the intermittent controller intervenes by replanning a movement often, then the horizon should be shortened as planning for longer movements is computationally costly [44]. Conversely, if the planned portion of the movement is executed without replanning, then the horizon should be lengthened so as to reduce the rate of replanning, thereby reducing the cognitive cost of the task.

When the whole task can be planned in one step because the uncertainty is minimal, then the horizon can be close to the duration of the movement. Importantly, as the preferred movement velocity can largely vary between individuals due to differences in individual vigor [45], the movement duration for a given task with minimal uncertainty may be determined by the concurrent minimization of effort and task completion duration [46, 47], resulting in a personalized optimal horizon. However, such minimization of task completion duration, which could be implemented in the cost function of Eq. 4, is out of scope of the present study.

Conversely, when there is uncertainty in the task due to additive motor noise [48], it will be desirable to reduce the planning horizon. This may also impact the duration of the whole movement, as additive motor noise may induce a bounded optimal movement time [49]. In Fig. 3b, we illustrate this effect of motor noise by computing the total cost of movement for a simple reaching movement with duration from 500 ms to 1.5 s to a static target. In such a case, the horizon can be chosen equal to the movement duration as the CNS can easily plan such simple reaching tasks at once if its estimation of **p**_*M*_ is accurate. In absence of noise (and any complementary costs related to time discussed above), the optimal horizon and movement duration would tend to infinity (Fig. 3b, right). However the optimal movement duration shortens as the level of motor noise increases, as evidenced in previous works [8].

**Figure 3:**
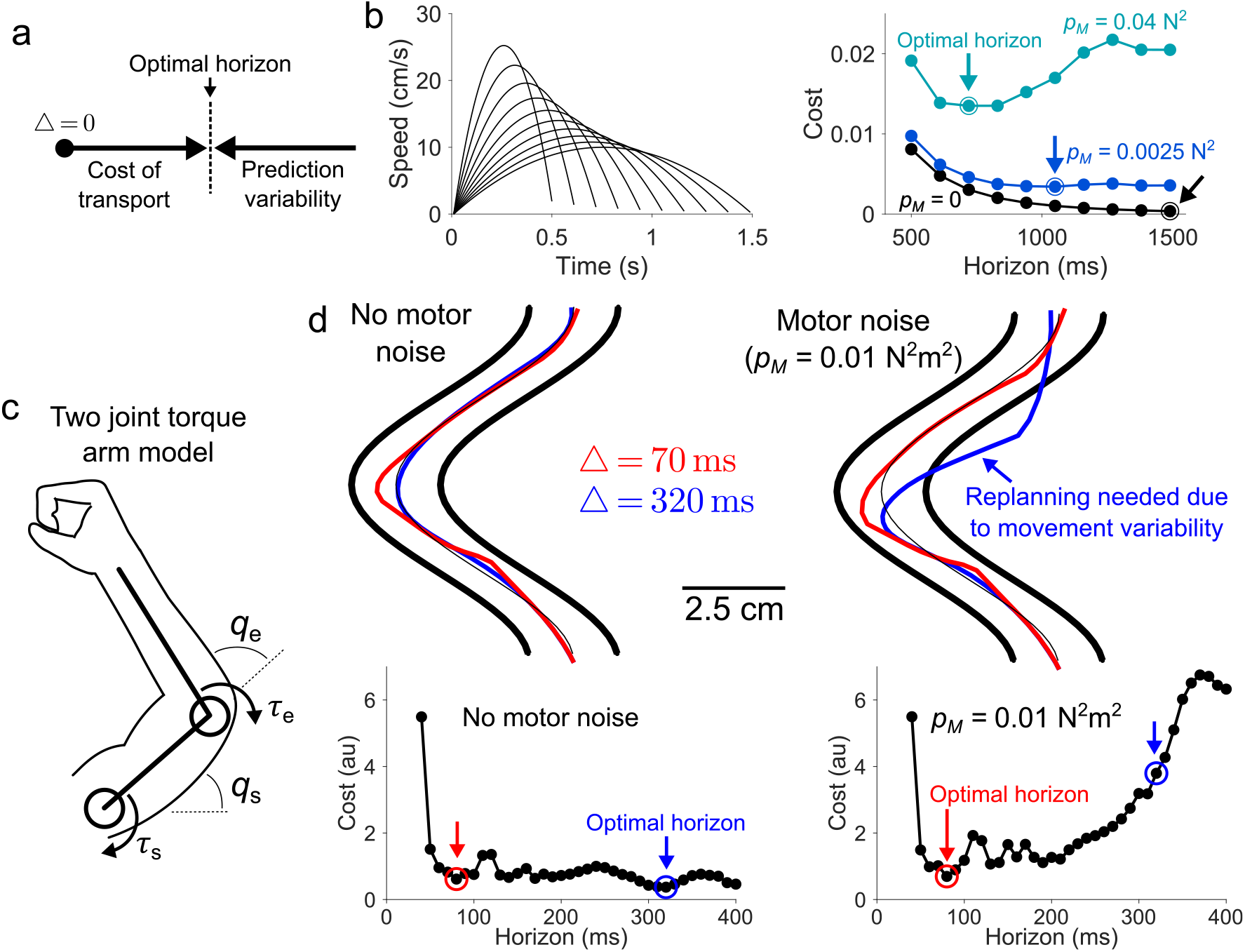
Optimal horizon depends on the cost of transport and prediction variability. (a) Minimizing the cost of transport leads to a longer horizon, but variability in the prediction error shortens it, yielding an optimal horizon. (b) Simulations of reaching for different horizons to the same target location 10 cm away (*p*_*S*_ = 0 and *p*_*T*_ = 100 cm^2^). As motor noise gradually increased (*p*_*M*_ = 0 N^2^ in black, *p*_*M*_ = 0.0025 N^2^ in blue, and *p*_*M*_ = 0.04 N^2^ in turquoise) the horizon yielding the smallest cost became shorter. (c) A two joint model of the arm with elbow and shoulder joints was used to simulate the effect of world model inaccuracy on the optimal horizon. (d) Intermittent controller with no motor noise (left) and motor noise *p*_*M*_ = 0.01 N^2^m^2^ (right) moving through a narrow path with walls (thick traces). With no motor variability (left), the cost was minimized by planning with a longer horizon of Δ= 320 ms (blue). A shorter horizon yielded a more intermittent movement that strayed from the center of the path (red). But when motor noise was introduced, a shorter horizon yielded a less costly movement overall relative to a longer horizon as the latter strayed greatly from the path due to an accumulation of error.

To further illustrate this effect, we simulated the effect of uncertainty when controlling a two-joint arm with elbow and shoulder torques (see Fig. 3c). In the simulation, the objective was to follow a target trajectory for 400 ms within a channel (Fig. 3d, top left). The horizon was defined as the amount of available information regarding the target trajectory, which depends on how long it is visible ahead of the current time. For example, a 40 ms horizon meant that only a tenth of the total target trajectory was visible ahead of the current time. Note that in this context, we assumed that the intermittent controller only replans a movement after fully executing its open-loop plan, meaning that FRC, PETC and IPC were similar. Uncertainty was introduced by simulating additive motor noise with variance *p*_*M*_ in the movement dynamics.

First, when *p*_*M*_ ≡ 0, a long horizon (Δ= 320 ms) yielded the optimal cost of movement (within the tested interval Δ ∈ [40, 400] ms) as the intermittent controller could plan most of the movement without generating significant task errors. This long horizon resulted in a smooth movement that optimally accounted for the target’s curvature (Fig. 3d, blue trajectory in top left). Conversely, with a shorter horizon (Δ= 70 ms) the tracking was less performant due to a reduced knowledge of the target’s curvature, resulting in overshoots and jerk at the junction of planned segments (Fig. 3d, red trajectory in top left).

However, a long horizon does not allow to complete the task successfully in presence of additive motor noise, as the error exceeds the task tolerance and the intermittent controller veered from the target trajectory, due to the time-accumulation of noise corrupting the plan’s execution (Fig. 3d, blue in top right). With the normal implementation of FRC, PETC and IPC, these errors would be corrected through replanning. Conversely, a shorter horizon (Δ = 70 ms) allowed to correct movement errors due to motor noise on-the-fly (Fig. 3d, red in top right).

These cases illustrate how the optimal movement duration and planning horizon will depend on the task parameters. In the end, one should aim at minimizing the amount of replanning steps (as this is costly [50]) and the overall cost of transport. This adaptation is consistent with the results of [30] in a similar task, where the horizon became longer for experts, suggesting that they could rely on a reduced motor variability [51], can leverage a longer horizon.

We therefore include the following adaptive mechanism aiming to minimize the number of replanning events in subsequent simulations:

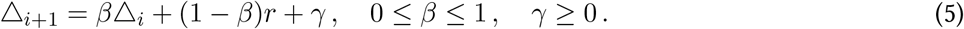

After each replanning, the next horizon Δ_*i*+1_ is adapted as a convex combination of the current horizon Δ_*i*_ and the time elapsed since the last replanning *r* depending on *β. γ* is a confidence parameter that tends to extend the horizon and guaranties Δ *>* 0 ms.

### 2.5 Human-like submovements in continuous tracking

In this section, we leverage simulations of a trajectory-tracking task with wrist flexion-extension to investigate the effect of different levels of sensory noise on the number of replanning events with our three models (Fig. S2a; see Methods for details) [52]. This allows to analyze the sensitivity of IPC to the third component of the admissible variance. FRC, PETC, IPC are used to track a moving target with low (*p*_*S*_ = 0.0025 cm^2^) and high (*p*_*S*_ = 2.25 cm^2^) sensory noise using an adaptive horizon with *β* = 0.4 and *γ* = 30 ms (Fig. 4a).

**Figure 4:**
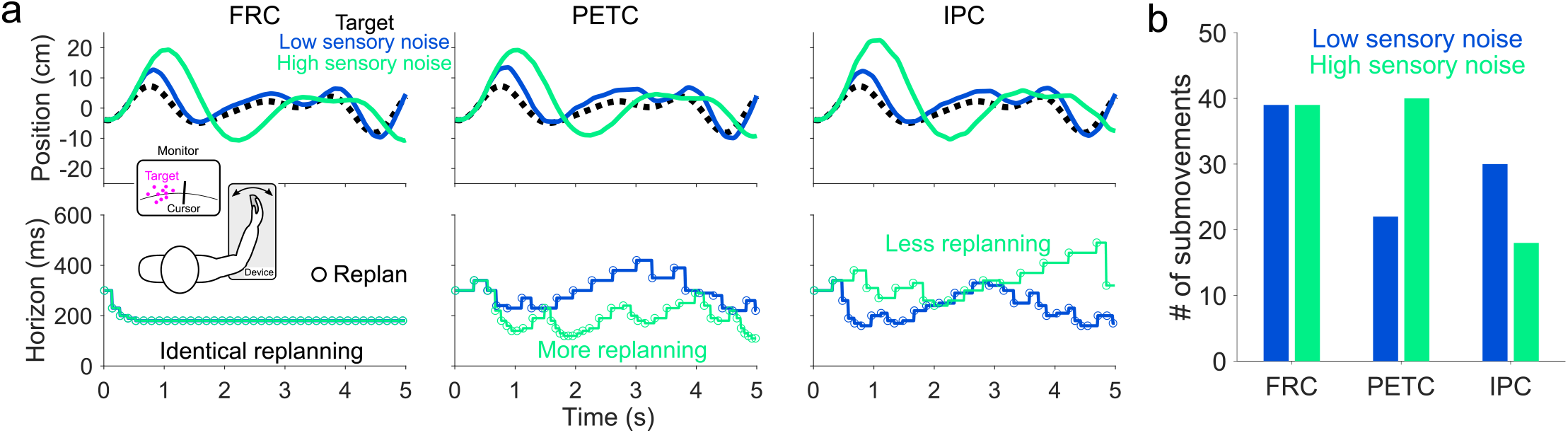
Only IPC predicts lower replanning rate when tracking a target under high sensory noise with an adaptive horizon. (a) Tracking using an adaptive horizon for low and high sensory noise using FRC, PETC, and IPC. Under FRC, horizon quickly adapts to the minimum value for both low and high sensory noise. With PETC, more replanning occurs with high sensory noise, leading to shorter horizons. IPC’s horizon is longer with higher sensory noise. (b) Number of submovements in (a) from each model separated by sensory noise level. FRC uses the same number of submovements irrespective of sensory noise level. PETC uses more submovements with greater sensory noise. Only IPC exhibits fewer submovements with high sensory noise.

As before with FRC, the number of submovements, defined as the number of replanning events during tracking, is similar for low and high sensory noise (Fig. 4b), as FRC replans at a constant rate, causing the horizon to plateau quickly to its minimum value. On the other hand, PETC has more submovements, and a shorter horizon on average, when tracking the same target with high sensory noise. The opposite is observed with IPC as it uses fewer submovements when the target is corrupted by high sensory noise. Thus, IPC’s horizon is larger with higher sensory noise. One of the main observations of the present paper is that only IPC allows to produce a decrease in the number of replanning events when increasing sensory noise, as reported in [29].

### 2.6 Replanning and optimizing a hunter’s movements

Finally, we investigate the effect of the target’s motion on the replanning rate of IPC allowing us to analyze the sensitivity of the intermittent controllers to the last component of the admissible variance: the target’s movement variability **p**_*G*_ = diag(**P**_*G*_). To do so, we simulate the scenario of a hunter tracking a prey’s movement in a plane (Fig. 5a). Specifically, we adjust the prey’s sway along the horizontal axis using a sine function with frequency *f* ∈ {0, 1, 3, 5} Hz, making the prey difficult to catch due to the increased goal variability *p*_*G*_ (Fig. 5b). The initial planning horizon is Δ_0_ = 400 ms and we use *β* = 0.4 and *γ* = 10 ms to adapt the horizon for all simulations.

**Figure 5:**
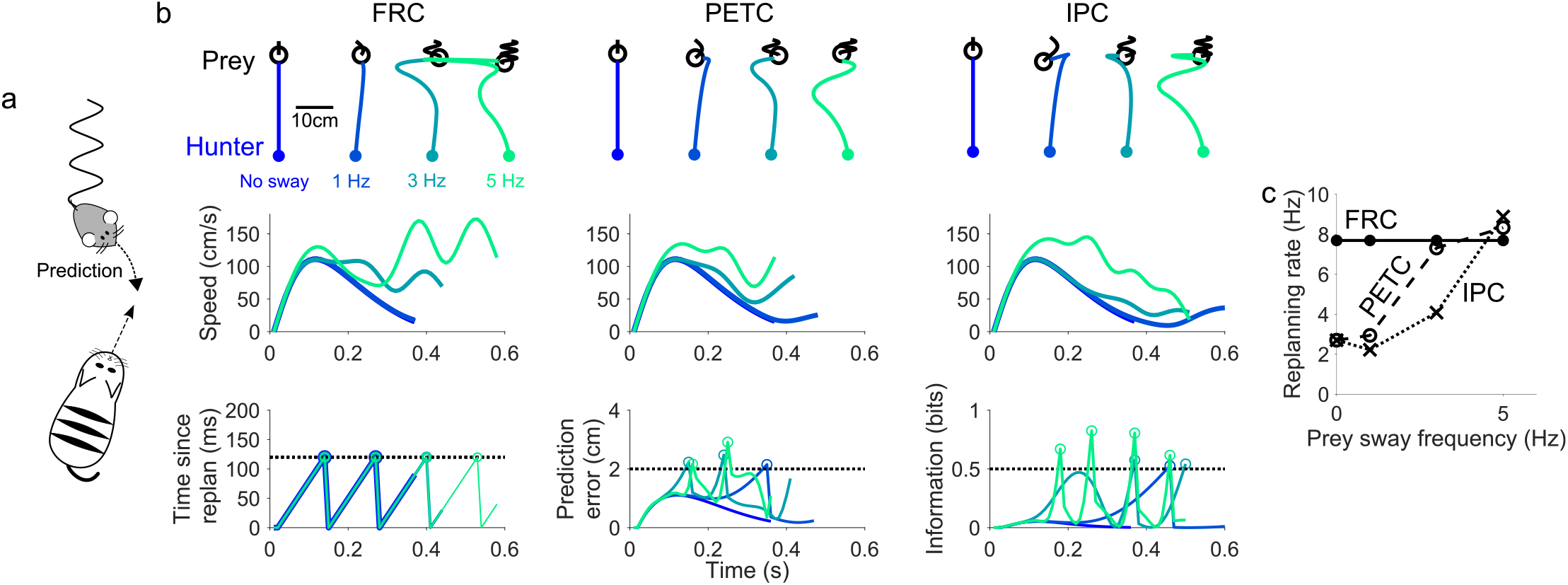
Simulation of a hunter intercepting a prey in a plane. Simulations were performed for different prey movement frequency. Note that the delay that sometimes seems to appear between the prediction error or information threshold crossing and the replanning event is only due to the simulation timestep of 10 ms. (a) Schematic of the simulation. FRC, PETC, and IPC were used to intercept a moving prey that swayed laterally according to a sinusoidal function. (b) Top panels show the hunter and prey’s trajectories are shown up until the time when the distance between the hunter and prey falls below *p*_*T*_. As the prey’s sway frequency increases, all intermittent controllers make lateral movements to anticipate the prey’s lateral motion. Greater sway causes the controllers to overshoot further along the lateral direction. Middle and bottom panels show the controller’s speed and replanning decisions as a function of time. The speed had more bumps as the prey’s sway frequency increased as the controllers made lateral adjustments. FRC replanned at a fixed interval irrespective of the prey’s sway frequency. The prediction error and information exceeded their thresholds for PETC and IPC more often as the prey’s sway frequency increased. (c) Replanning rate as a function of the prey’s sway frequency. The replanning rate was constant for FRC, but it increased with frequency for both PETC and IPC.

FRC, PETC, and IPC made lateral movements to intercept the prey as it swayed laterally (Fig. 5b, top panels). This could be seen by the increasing number of bumps in the speed time-series (Fig. 5b, middle panels). As the prey’s sway frequency increased, both the prediction error and information exceeded their respective thresholds more often due to uncertainty in the prey’s lateral movements (Fig. 5c, bottom panels). FRC’s replanning rate remained constant for all sway frequencies, but PETC and IPC’s replanning rate became larger as the prey’s sway frequency increased (Fig. 5c).

These simulations show how the goal variability changes the behavior of the intermittent controllers. We have now looked at how the sensory noise, task tolerance, motor noise and goal variability each affect IPC’s replanning behavior independently of each other. We can now look at how these four variables interact with one another by varying them together. We varied the sensory noise *p*_*S*_ ∈ {0.01, 1, 9, 100} cm^2^, task tolerance *p*_*T*_ ∈ {1, 4, 16, 25} cm^2^, and motor noise *p*_*M*_ ∈ *{*0, 4, 16, 100} N^2^m^2^ concomitantly in a 4D parameter space to examine the interactions between the four parameters determining IPC’s behavior (Fig. 6). Their effect on IPC was examined through IPC’s replanning rate, horizon, and path length when intercepting the prey.

**Figure 6:**
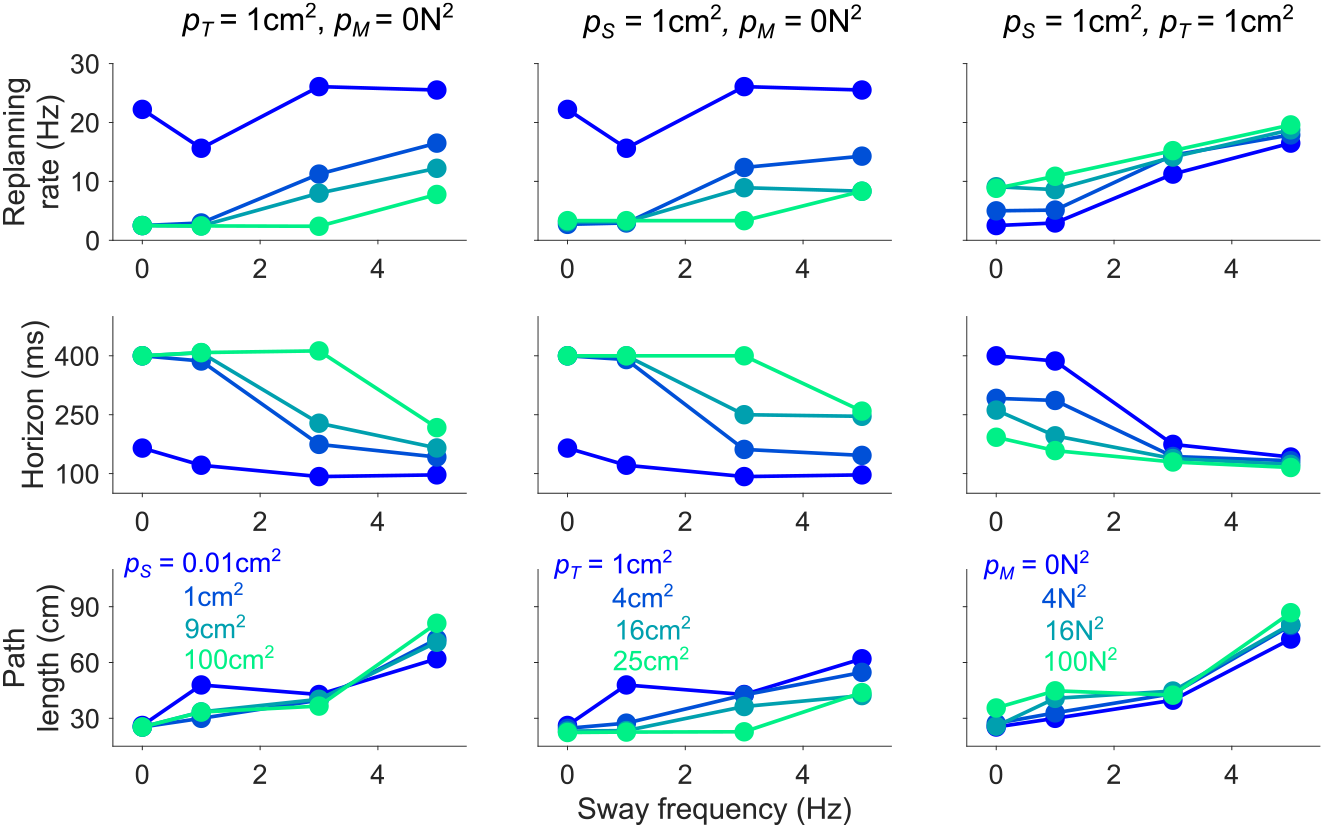
Sensitivity of IPC to the parameters determing *p*_*e*_. Simulations were performed with IPC where the prey’s sway frequency, and estimated sensory noise *p*_*S*_, task tolerance *p*_*T*_, and motor noise *p*_*M*_ were varied independently. Replanning rate, mean horizon during the task, and the total path length of IPC as a function of sway frequency for varying values of sensory noise *p*_*S*_ (left column), task tolerance *p*_*T*_ (middle), and motor noise *p*_*M*_ (right). Replanning rate increased with sway frequency (top panels) but larger sensory noise and task tolerance decreased the overall rate. Greater noise, on the other hand, increased replanning rate throughout. The horizon exhibited the opposite trends relative to the replanning rate as the horizon adapted to be shorter as replanning increased and vice versa.

Fig. 6 shows IPC’s replanning rate, horizon, and path length as a function of the prey’s sway frequency for different values of sensory noise *p*_*S*_ (left column), task tolerance *p*_*T*_ (middle column), and motor noise *p*_*M*_ (right column). The replanning rate decreases with increasing sensory noise and task tolerance but increases with motor noise. Conversely, the adaptive horizon increases with increasing sensory noise and decreases with motor noise, as implemented by Eq. 5. A higher replanning rate allows IPC to update its motion plan more often, leading to a shorter path length when intercepting the prey, with the exception when the task tolerance is changed because a larger sized prey can be intercepted more easily, thus the path length becomes considerably shorter as the task tolerance and prey size increase.

## 3 Discussion

The Information Predictive Control introduced in this paper is a computational framework to explain how the brain plans and executes movements under uncertainty and time-varying goals. IPC integrates an information module with model predictive control [32, 53] enabling replanning when deviations from expected states become sufficiently surprising, i.e., when they contain sufficient information. By combining control and information theory, IPC addresses the limitations of previous models of intermittency that did not consider how task tolerance, sensorimotor noise, and goal variability influence the rate of replanning [21, 25–28]. In the present paper we analyzed the main features of IPC: (i) replanning based on a surprise threshold accounting for the task tolerance, sensorimotor noise, and task uncertainty, and (ii) a planning horizon adaptation mechanism minimizing the cost of replanning. As other discrete models [25, 28], IPC explains and reproduces century-old observations of intermittency in movement control, which were reported in discrete [22, 54] and continuous human movements [55, 56], and in other species such as cats [19], mice [20], and non-human primates [24]. This intermittency has been proposed to stem from a brain sensitive to errors only above a certain threshold [28, 57, 58] or be triggered by alpha waves at a fixed rate [25]. However, these hypotheses did not account for the influence of sensory, motor, and goal noise on the replanning rate.

In the IPC framework, when the prediction error exceeds its permissible variance, the generated information triggers movement replanning. Since replanning a movement entails metabolic costs [50], cognitive costs [59], and can impair motor performance [60], IPC minimizes the replanning rate by considering several factors. First, we have shown that, when compared to the FRC model of intermittent control [25, 28], PETC and IPC lead to a lower number of replanning events when reaching in an unknown force-field, resulting in similar predictions when there is no sensorimotor noise or uncertainty in the task. Second, it can be reasonably assumed that large sensory noise in vision or proprioception increases the permissible variance in the prediction error, as the error estimates become less reliable. In the present paper, we have shown that IPC reduces the replanning rate when sensory information is noisy, consistent with the findings in [29] that intermittency decreases when tracking a target with noisy peripheral visual feedback. In contrast, with FRC the number of replanning events remained constant, due to the constant task duration, and with PETC the number of replanning events increased, due to the constant prediction error threshold being exceeded more often in less reliable environments. Third, in IPC the sum of motor and goal variability in the permissible variance is capped by the task tolerance, i.e. the accuracy constraints of the task. Greater motor or goal variability will reduce the replanning frequency as long as the task tolerance is high, meaning that large deviations along task-irrelevant directions are neglected [13,36], while deviations along task-relevant directions are tolerated only if they do not impact task success. Consequently, if the task tolerance is low, or with high accuracy constraints, larger motor or goal variability raises the replanning frequency, as even small deviations in the prediction error trigger replanning. This minimum intervention principle has been described through the concept of the uncontrolled manifold, according to which only degrees of freedom (DoF) relevant to the task are controlled [61,62]. While OFC also avoids intervening along task-irrelevant DoFs, and can adapt feedback gains to changing constraints in task goals with appropriate modifications in the simulated strategy [15,17,63], it intervenes along the task-relevant dimension, regardless of the error magnitude or target reachability. In contrast, IPC intervenes only when the error is substantial enough to warrant replanning and the target is likely reachable. IPC thus extends and refines the minimum intervention principle by distinguishing intervention needs along task-relevant as well as task-irrelevant dimensions.

Our simulations show how the planning horizon—the duration into the future considered when planning a movement—emerges as a property tied to the replanning rate. When only the cost of movement was considered in IPC, the optimal horizon extended toward infinity as slower movements were least costly. In practice, movement duration is limited, which can be modeled via a reward discounted with time yielding an optimal movement duration [64, 65]. Experimental evidence also suggests that very slow movements may be as costly for the brain as very fast, effortful, ones [47, 66]. In our approach, uncertainty in the prediction error naturally limits the planning horizon, consistent with variance-minimization frameworks [8, 49].

Throughout the paper, we showed how inaccuracies in the environment and task model promote a shorter planning horizon, as frequent replanning reduces the overall task cost. This prediction aligns with findings from a study in which participants had to move a cursor along a narrow path [30]. Novice participants employed a short horizon, likely due to the need for frequent error corrections. In contrast, experienced participants used a longer horizon, possibly by leveraging task knowledge acquired through practice that reduced execution errors and replanning needs. The adaptive mechanism proposed in the present paper similarly increased the horizon in predictable conditions and reduced it when task uncertainty increased, as illustrated in the hunting example. This mechanism could explain why movement intermittency increases and the duration of submovements decreases when approaching a small target [67].

While we used a surprise threshold to trigger replanning, our approach could be extended by incorporating principles of decision-making. For instance, a drift-diffusion model could trigger replanning based on accumulated information over time [68], which could make IPC more robust to false alarms and enable it to correct permissible but persistent prediction errors.

A significant advantage of IPC is the flexibility of the underlying internal model, which can be linear or non-linear, take any form from differential equations to neural networks, as long as it can predict the future internal and environment’s state. This work focused on IPC’s properties, but an internal model could be integrated and learned through feedback error minimization [35, 39] or reinforcement learning [69]. As the internal model better compensates for the environment dynamics, it will yield more accurate predictions, thereby reducing replanning events. Furthermore, IPC’s flexibility in modeling the cost minimized by the brain can incorporate factors such as the movement timing and reward discussed above, or feedback mechanisms such as short and long-delay reflexes. Importantly, IPC’s cost function could also consider system stochasticity to predict key motor control features like feedforward control of motion [7, 22] and cocontraction [9, 10]. Finally, the fact that IPC only replans movements when a surprise threshold is exceeded does not imply that it cannot predict early corrective forces observed when a mechanical perturbation is applied [16, 70]. In fact, considering a stochastic cost function accounting for noise and uncertainty, as in [9, 10], and reflexes, which can also be impacted by sensory information [41], is feasible and would predict such forces.

Overall, IPC’s ability to produce and correct movement trajectories based on an internal model, information theory, and a flexible formulation of the brain’s internal costs, make it a useful tool for computational neuroscientists to emulate and investigate how the brain’s control movements in a dynamic environment.

## 4 Methods and Materials

The intermittent FRC, PETC, and IPC use an internal model to plan the future control inputs **u** needed to accomplish a given goal over a *time horizon* Δ. Here we describe how each model works on the simulations described in the paper.

### Intermittent control of a linear system

We first simulate reaching movements using an arm modeled as a point mass in two dimensions *x* and *y* (as illustrated in Fig. 2). The arm moves according to the linear dynamics

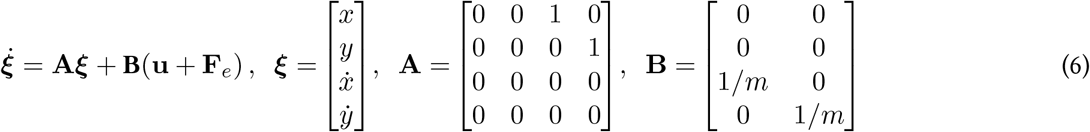

where ***ξ*, u, F**_*e*_ are functions of time *t* and *m* = 1 kg. **F**_*e*_ is the force from the environment, which is **0** in the free condition, or

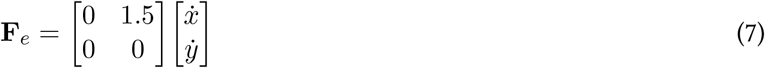

as a lateral force field, where **F**_*e*_ is in N and 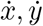 in m*/*s. The goal is to move state ***ξ***(*t*) at current time *t* to the goal ***ξ***^∗^(*t*^+^) at *t*^+^ ≡ *t* + Δ using a force **u**(*ζ*) =[*u*_*x*_(*?*), *u*_*y*_(*?*)]′, *ζ*∈ [*t, t*^+^], by minimizing the cost function

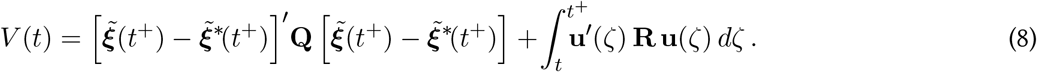

The first term weighted by **Q** is to ensure that the predicted state 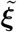 comes close to the predicted goal 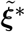 at the horizon, the second term to minimize effort.

The future state 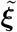 is computed by integrating Eq. 6. The future 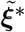 is estimated using linear quadratic estimation [71], starting from their last measured value at time *t*. For this purpose, the goal state is assumed to evolve according to 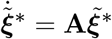 and observed with

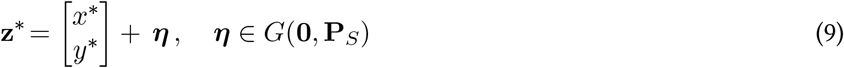

where **P**_*S*_ is the sensory noise covariance matrix.

The future control inputs **u** to minimize the cost of Eq. 8 were optimized by Quadratic Programming (QP), where the control inputs were iteratively updated until the solution converged to an acceptable minimum. There are several algorithms for solving QP class problems, and the one we adopted was the interior-point method implemented in Matlab’s ‘*fmincon*’ function due to its speed and robustness. The last control input was bounded to be equal to **0** to avoid discontinuities between movements and to keep IPC passive when time exceeded the horizon.

Above equations were discretized with a 1 ms time step for all simulations. In the simulations of Fig. 2a (solid blue and dashed blue), reaching movements were simulated from the origin **x**_0_ = [0, 0]′ toward a target located at [0, 0.1] m using *p*_*S*_ = 0.01 cm^2^, Δ ∈ {500, 1000} ms, **Q** = diag(30, 30, 1, 1) and **R** = 10^−5^𝕀_2_, where 𝕀_2_ is the 2×2 identity matrix. For PETC, we varied the prediction error threshold to be {0.5, 0.7, 1, 2} cm. For IPC, the target tolerance was *p*_*T*_ = {0.25, 0.49, 1, 4} cm^2^ to observe its change in replanning rate. For Fig. 3b we simulated 30 reaching movements with the same parameters as above while the horizon Δ was varied between [500, 1490] ms in steps of 110 ms. For these simulations, constant motor noise was added to the motor command: **u** + ***ν, ν*** ∈ *G*(**0**, *p*_*M*_ 𝕀_2_). For each value of Δ, movements were simulated with *p*_*M*_ = 0 N^2^, *p*_*M*_ = 0.0025 N^2^ and with *p*_*M*_ = 0.4 N^2^.

For the simulations in Fig. 5b, the horizon was initially set to Δ= 400 ms then adapted according to Eq. 5 with *β* = 0.4 and *γ* = 10 ms. The other parameters were *p*_*S*_ = 4 cm^2^, *p*_*T*_ = 4 cm^2^, *p*_*M*_ = 0, **Q** = diag(10^3^, 10^3^, 10, 10) and **R** = 10^−1^𝕀_2_. The prey moved according to

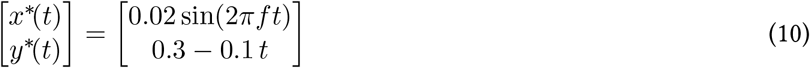

where the frequency *f* was selected from the set {0, 1, 3, 5} Hz.

To examine IPC’s sensitivity to its parameter settings when intercepting the prey (Fig. 6), we varied the sensory noise *p*_*S*_ ∈ {0.01, 1, 9, 100} cm^2^, task tolerance *p*_*T*_ ∈ {1, 4, 16, 25} cm^2^, and motor noise *p*_*M*_ ∈ {0, 4, 16, 100} N^2^m^2^ independently of each other and calculated IPC’s replanning rate, average horizon, and path length defined as the total Euclidean distance covered by IPC until interception.

### Simulation of wrist tracking movement

For the tracking simulations in Fig. 4 and Fig. S2, the target was composed of a multi-sine function

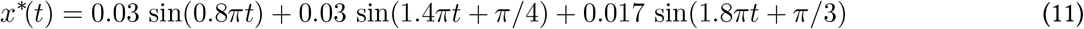

where *t* is the time. FRC, PETC, and IPC tracked this target for five seconds. Each intermittent controller tracked the target movement along the single *x* dimension using the cost function

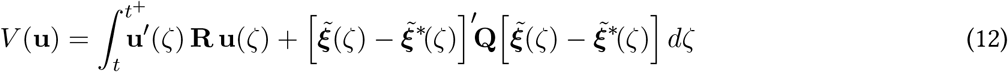

with *p*_*T*_ = 0.25 cm^2^, **Q** = diag(20, 20, 0.1, 0.1) and **R** = 10^−8^ 𝕀_2_. In the simulation with the adaptive horizon (Fig. 4), the initial horizon was set to Δ = 300 ms with *β* = 0.4 and *γ* = 30 ms. Low and high sensory noise were simulated with *p*_*S*_ = 0.0025 cm^2^ and *p*_*S*_ = 2.25 cm^2^, respectively. The task tolerance was *p*_*T*_ = 1 cm^2^ and the prediction error threshold was 1 cm.

In the sensitivity analysis (Fig. S2), the horizon was constant throughout each trial but was selected to be Δ ∈ {200, 400, 600, 800} ms. PETC was simulated with prediction error thresholds of {1, 5, 10, 20, 30} cm and IPC was simulated with *p*_*S*_ ∈ {0.0025, 2.25, 25, 100, 225} cm^2^ to examine the sensitivity of their replanning rates to these parameters.

### Simulation of two-joint arm control

The effect of model inaccuracy on IPC’s behavior was simulated in Fig. 3d using a model horizontal arm movement around the shoulder and elbow with joint angles **q** = [*q*_*s*_, *q*_*e*_]′ controlled by the torques ***τ*** = [*τ*_*s*_, *τ*_*e*_]′ (Fig. 3c), having dynamics

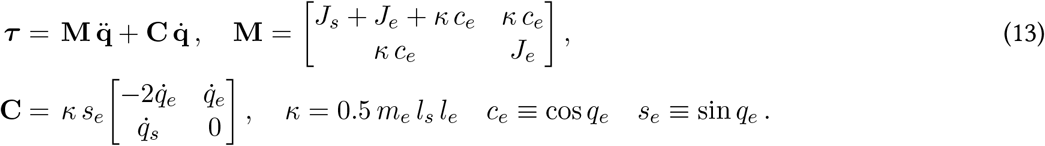

We used the parameters from [71]: *l*_*s*_ = 0.3 m and *l*_*e*_ = 0.35 m for the lengths of the shoulder and elbow segments, *J*_*s*_ = 0.05 kg m^2^ and *J*_*e*_ = 0.06 kg m^2^ their moments of inertia, and *m*_*s*_ = 1.9 kg and *m*_*e*_ = 1.7 kg their masses. The joint inertia was computed assuming that the elbow’s center of mass is in the middle of its segment.

The state 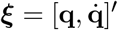 was controlled to track the Cartesian goal trajectory in the middle of the path

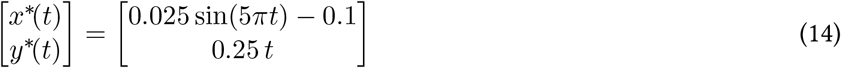

which was converted to joint coordinates via the arm’s inverse kinematics

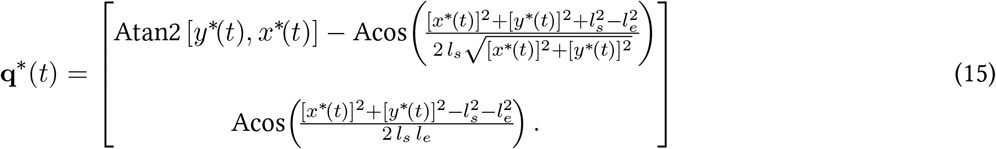

to yield 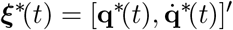. This goal state was tracked by minimizing cost function Eq. 12 with **Q** = diag(10^3^, 10^−2^) and **R** = 10^−3^ 𝕀 _2_. The horizon was incrementally changed from 40 ms to 400 ms in steps of 10 ms. The prediction error was set to a large value *p*_*e*_ = 100 cm^2^ so that movement replanning was suppressed in these simulations.

## Acknowledgments

The authors would like to thank Yasuharu Koike, Hiroaki Gomi and Frédéric Crevecoeur for discussions on this paper’s material.

## Materials & Correspondence

Correspondence and requests for material should be sent to A.T.

## Data availability

The datasets analyzed in the current study are available in the Figshare repository (https://doi.org/10.6084/m9.figshare.28586249).

## Supplementary Materials

### 4.1 Reaching in a force field

FRC, PETC, and IPC can reproduce human-like behavior when reaching in a velocity-dependent force field that applied an external force

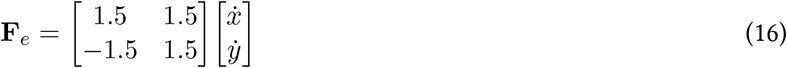

that pushed the controller in a clockwise direction. We simulated FRC, PETC, and IPC reaching under four conditions: the null field condition with **F**_*e*_ = **0**, a force field condition where the controller was unaware of the force field, a force field condition after the controller was given full knowledge of the force field dynamics, and an after-effect condition where **F**_*e*_ = **0** with the controller still under the belief that the force field was active. All controllers were simulated with Δ = 500 ms, *p*_*S*_ = 0.01 cm^2^, **Q** = diag(30,30,1,1), and **R** = 10^−5^𝕀 _2_. PETC’s prediction error threshold was 1 cm and IPC’s task tolerance was set to *p*_*T*_ = 1 cm^2^.

Fig. S1 shows each controller’s trajectories as it reached eight equidistant targets placed 10 cm from the origin. In the null field condition, all controllers used a straight movement to reach each target (Fig. S1, left). When the force field was imposed without the controller’s knowledge, the force field pushed the controller in the clockwise direction, causing all of its trajectories to curve in that direction (Fig. S1, middle-left). Once the intermittent controller was given full knowledge of the force field’s dynamics, it was able to reach the targets with near-straight trajectories (Fig. S1, middle-right). Finally, once the force field was removed without the controller’s knowledge, it overcompensated in the counter-clockwise direction, leading to an after-effect commonly observed after human motor learning (Fig. S1, right).

**Figure S1:**
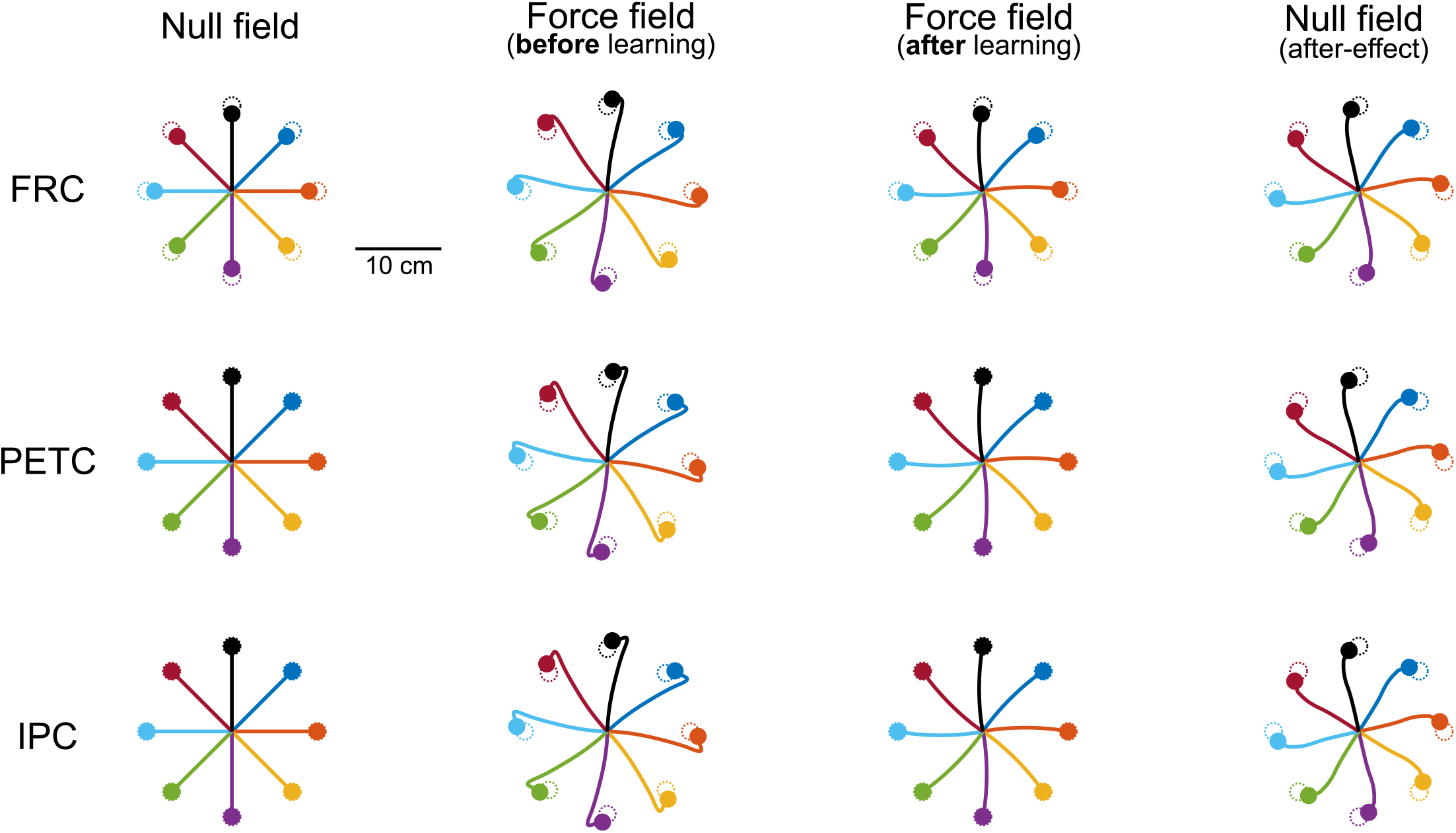
FRC, PETC, and IPC reproduce human-like behaviour when learning to reach in a velocity-dependent force field. In the null field, all controllers move straight towards the target (left). When a velocity-dependent force field is imposed without the controller’s knowledge, its trajectory curves in the clockwise direction (middle-left). Once the controller is given full knowledge of the force field, it recovers near-straight reaching trajectories (middle-right). When the force field is abruptly removed without the intermittent controller’s knowledge, IPC exhibits an after-effect by overcompensating and curving its trajectory in the counter-clockwise direction (right).

### 4.2 Tracking with a constant horizon

We simulated tracking in one dimension with a constant horizon using FRC, PETC, and IPC to illustrate how the replanning rate changes with different levels of sensory noise. Fig. S2b shows the trajectories of the target and FRC, PETC, IPC tracking with small and large sensory noise and a constant horizon Δ = 200 ms. FRC’s replanning rate is again independent of the sensory noise. For PETC, we tested different prediction error thresholds to observe changes in its replanning rate (Fig. S2c). These simulations were carried out for four constant horizons Δ = {200, 400, 600, 800} ms. While PETC’s replanning rate decreases as the prediction error threshold increases, the replanning rate with high sensory noise is always larger or equal to the low sensory noise simulation. Hence, PETC uses a larger replanning rate when there is higher sensory noise of the target’s position.

We then carried out simulations with IPC with different values of *p*_*S*_ to examine the effect of sensory noise on IPC’s replanning rate (Fig. S2d). Like with the PETC simulation, four constant horizons were simulated. As sensory noise *p*_*S*_ increased, IPC’s replanning rate decreased. IPC’s replanning rate was always lower with high sensory noise relative to the low sensory noise simulation.

**Figure S2:**
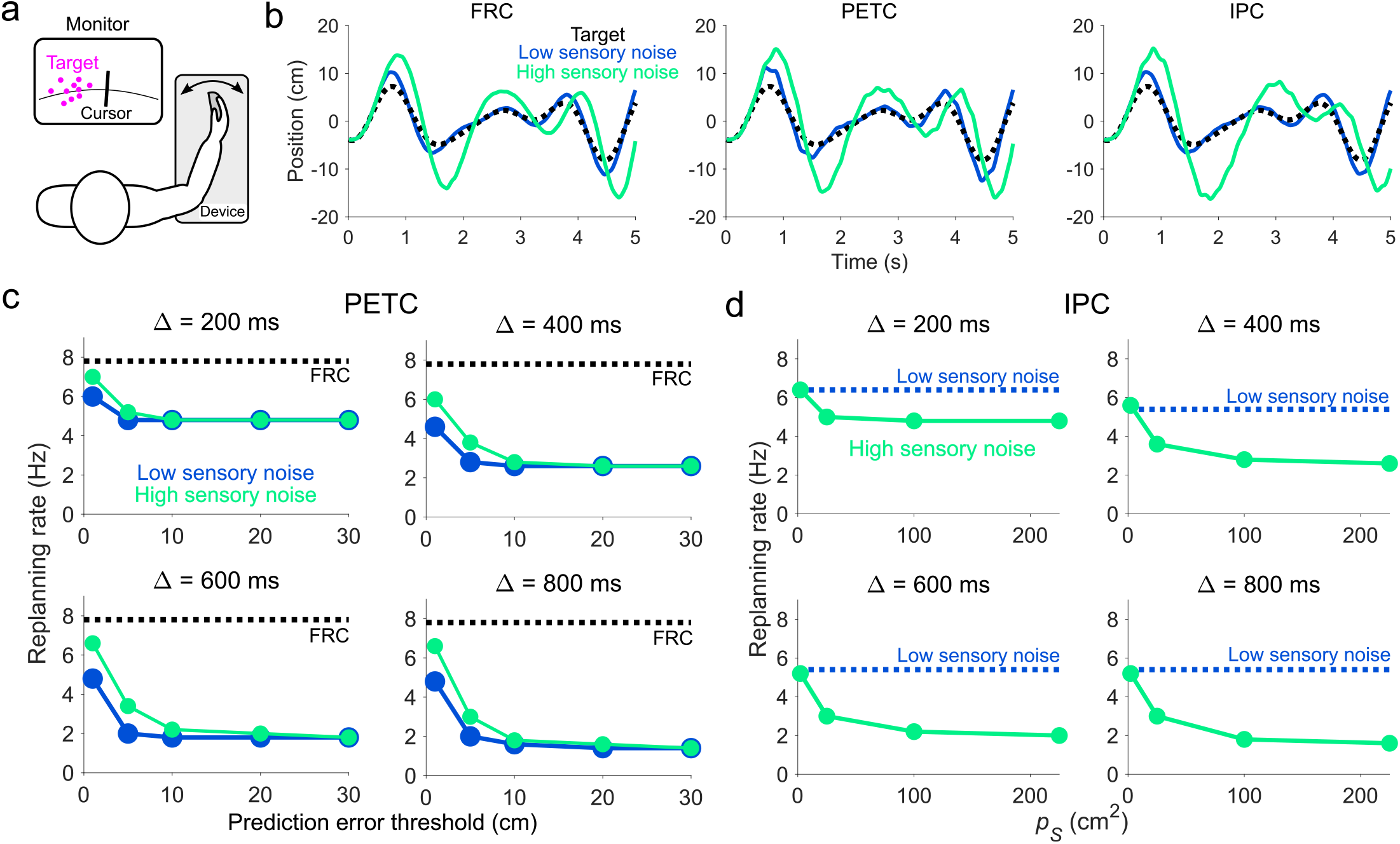
IPC’s replanning rate decreases with larger sensory noise relative to other models during tracking. (a) Illustration of a one-dimensional tracking task with visual feedback or sensory noise. (b) Trajectories from FRC, PETC, and IPC when tracking a target (black) with low (blue) and high sensory noise (green). (c) PETC’s replanning rate as a function of the prediction error threshold for low and high sensory noise. Replanning rate with high sensory noise was always greater or equal to the replanning rate with low sensory noise. For FRC, the replanning rate was constant regardless of sensory noise level (dashed black line). (d) IPC’s replanning rate during tracking as a function of *p*_*S*_. Relative to the replanning rate for small sensory noise (dashed blue line), replanning rate decreases as *p*_*S*_ increases for all horizon values.

